# Decision Voting Based Multiscale Convolutional Learning of Brain Networks With Explainability

**DOI:** 10.1101/2025.10.29.685275

**Authors:** Deep Sekhar Ghosh, Sanjay Ghosh

## Abstract

The diagnosis of neurological disorders requires comprehensive frameworks that incorporate multimodal neuroimaging data while ensuring clinical interpretability. Recent neuroimaging research is focusing on the integration of brain structure and function to reveal some of the prominent alteration caused by a brain disorder at the system level. This work presents a fresh multiscale graph convolutional network (GCN) framework that integrates structural connectivity (SC) from diffusion tensor imaging and functional connectivity (FC) from resting-state fMRI on three different anatomical scales. We present softmax-based decision fusion for cross-modal multiscale integration in our architecture. The preprocessing pipeline improves connectivity representations by means of graph diffusion, topological sparsification, and noise augmentation. Using five-fold cross-valuation, evaluated on a schizophrenia classification dataset, our model achieves 71.59% accuracy, outperforming single-scale methods and conventional machine learning benchmarks. Explainability analysis reveals that different dysconnectivity patterns in schizophrenia patients overlapping with biomarkers reported in literature. The multiscale approach shows complementing insights: coarse scales capture global network changes while finer scales identify localized subcortical disruptions. Combining diagnostic precision with biologically interpretable modeling, this work creates a new paradigm for interpretable brain network analysis.

## 1. INTRODUCTION

**N**EUROLOGICAL disorders, including schizophrenia, Alzheimer’s disease, Parkinson’s disease, epilepsy, autism spectrum disorder (ASD), major depressive disorder, schizophrenia, sclerosis etc. represent significant public health challenges [1]–[3]. These disorders exhibit their complex effects on brain connectivity networks [4]. Typically resulting in significant disruptions in daily life, these diseases affect cognitive, emotional, and motor skills. Recent advances in neuroimaging techniques have given insights into the structural and functional aspects of brain connectivity, helping researchers to model neurological disorders as disruptions in the brain’s complex networked architecture [5], [6]. Alteration in brain connectivity associated with mental disorders have been widely reported in both functional MRI (fMRI) and diffusion MRI (dMRI). Construction of brain connectivity/graph generally rely on a specific brain parcellation to define regions-of-interest (ROIs) within 3D volume of brain [7]. In the brain graph, nodes represent cortical and subcortical gray matter volumes and edges stand for the strength of structural or functional connectivity. Tractography algorithms applied to diffusion magnetic resonance imaging (MRI) or diffusion tensor imaging (DTI) data to estimate structional connectivity. Pairwise correlation between activation signals in various brain regions is measured by various functional brain imaging modalities - functional MRI (fMRI) is referred as functional connectivity. Graph Neural Networks (GNNs) have emerged as a powerful tool for studying brain connectivity to predict neuropsychiatric disorders associated with abnormalities in the functional and/or structural connectivity networks of the brain [8]–[10]. For instance, Ghosh et al. [11] applied graph convolutional neural networks (GCNNs) for schizophrenia classification using multimodal brain connectome data, achieving superior performance over traditional machine learning methods. Similarly, graph-based methods have provided valuable insights into other neurological conditions, including Alzheimer’s disease [12], autism spectrum disorder (ASD), and epilepsy [13]. Yao et al. [14] proposed a triplet graph convolutional network to combine coarse-to-fine parcellations of brain regions for ADHD classification.

Multimodal integration of functional connectivity (FC) derived from resting-state fMRI and structural connectivity (SC) derived from diffusion tensor imaging (DTI) offers complementary insights into the temporal and anatomical dynamics of the brain [15]. However, few studies leverage the full potential of combining structural and functional modalities to uncover overlapping and scale-specific characteristics of brain disorders. Unlike a single spatial scale brain graph, there could be partition each brain into multiple ROIs for generating multi-scale brain functional/structural connectivity networks, with each template corresponding to a specific spatial scale and ROI definition. Very limited attentions are made so far in simultaneously using multi-scale brain graphs for classification of brain disorders. Authors in [16] proposed to build on the stacked layers of graph convolution and the atlas-guided pooling, for a comprehensive extraction of diagnostic information from multiscale fully connected layers. In [14], a template matching-based triplet GCN (TGCN) model was proposed to learn functional / structural representations of brain connectivity networks at each scale. A multi-scale pooling was introduced to obtain representations of brain connectome at various scales [17]. These limited yet promising performance of multiscale GCN methods motivate us to pursue dedicated research towards a light-weight, simple, and efficient method for graph convolutional learning for brain networks. In this work, we introduce a new multi-scale (and multi-modal) framework to model brain connectivity networks,addressing these challenges through significant contributions. We employ spectral graph convolutions with a Chebyshev polynomial approximation to hierarchically learn connectivity features at multiple scales (83, 129, and 234 ROIs). Then we introduce a softmax-based decision fusion mechanism to dynamically integrate information across scales and modalities, optimizing classification performance. Finally, we incorporate GNNExplainer to provide explainability, revealing distinct biomarkers and connectivity patterns that underpin our classification decisions. Applied to the task of classification of schizophrenia, our framework achieves high diagnostic accuracy and identifies biologically interpretable connectivity disruptions, bridging the gap between advanced computational techniques and clinical applications in neuroscience.

The contributions in this work are summarized as follows.

### 1) Multiscale learning of brain networks

To the best of our knowledge, our study is the first to incorporate multiscaling brain connectivity networks in graph convolutional neural networks within a unified framework. We proposed a novel multiscale graph learning deep network for classifying brain disorders from brain connectivity map constructed on different scales: 83, 129, and 234. Our multiscale approach integrates brain connectivity data across different parcellations to capture both local and global interaction patterns.

### 2) Decision voting scheme

We introduce a data-driven fusion scheme referred as *decision voting* for integrating multiscale (and multimodal) features within our deep learning framework. Unlike concatenation-based fusion, our method predominantly learns the weights from the input data to combine the convoluted intermediate feature, resulting in superior performance.

### 3) Explainability analysis

To provide insights into which regions and connections of the brain are most influential in distinguishing between diseased patients and healthy controls, we perform an elaborate explainability analysis of our proposed method. Experimental findings suggest a complex pattern of altered connectivity in schizophrenia, involving frontal, temporal, and parietal regions.

### 4) Improved classification of brain disorder

The experimental results on schizophrenia classification demonstrate that our proposed *decision voting GCN* (DV-GCN) achieves 71.59% accuracy, outperforming single-scale methods and conventional machine learning benchmark.

Rest of the paper is summarized as follows. In Section I, we first discuss a quick summary of the recent research in brain connectivity based disorder classification using graph convolutional neural networks. This is followed by literature on explainability analysis in medical imaging. Section III covers the datasets and pre-processing pipeline. In Section III, we present our multiscale graph learning method in detail. Section V presents the experimental results on schizophrenia classification and the comparison with single-scale methods and conventional machine learning benchmark. We then also analyze explainability by identifying the influential brain areas that contribute to classification. Finally, we conclude the paper with potential future research direction in Section VI.

## RELATED WORK

Recent advances in multimodal and multiscale brain network analysis using graph convolutional neural networks have demonstrated promising results for neuropsychiatric disorder classification.

### A. Graph Learning in Brain Disorder Classification

Graph convolutional neural networks is found to achieve strong performance in neuroimage analysis [18], [19]. The benchmark for brain network analysis with graph neural networks provided a compact review of geometric deep learning for for neuroimaging analysis. Another benchmark study in [19] presented the state-of-the-art graph machine learning in brain connectomics. In one of the first research on deep learning of brain graphs, Kawahara et al. [20] introduced a new convolutional deep learning method for leveraging topological locality in brain networks. The method could also predict neurodevelopmental outcomes in preterm infants. A metric learning method with spectral graph convolutions on brain connectivity networks was presented in [21]. A local-to-global GNN (LG-GNN) with a local ROI-GNN for biomarker identification and a global Subject-GNN for inter-subject relationships was introduced in [6] presented. The ROI-GNN employs self-attention pooling to retain discriminative brain regions, while the subject-GNN uses adaptive weight aggregation for feature learning for for ASD and Alzheimer’s classification.

A hierarchical graph convolutional network for brain disorder diagnosis using functional connectivity networks derived from resting-state fMRI was proposed in [16]. A key innovation is the introduction of atlas-guided pooling, which leverages biologically meaningful hierarchical relationships among brain regions to perform nodal aggregation across scales [16]. By stacking graph convolution and atlas-guided pooling layers, the model comprehensively extracts diagnostic features from multiscale connectivity, leading to superior performance and interpretability compared to single-scale and data-driven pooling methods. Song et al. [22] applied hypergraph signal processing techniques to unravel higher-order brain network alterations in schizophrenia patients, further advancing the understanding of the disorder. Kong et al. [17] proposed a multi-connectivity representation learning network including structural, static, and dynamic functional connectivities for MDD diagnosis. The approach presents Static-Dynamic Fusion (SDF) modules to spread stationary connectivity patterns to dynamic graphs via attention mechanisms and Structural-Functional Fusion (SFF) modules to decouple modality-specific and shared features. A Multiview graph convolutional networks (GCNs) are being used to analyze neuroimaging data in Parkinson’s disease (PD) for improved diagnosis and understand disease progression was proposed in [23]. Liu et al. [24] developed a spatio-temporal hybrid attentive graph network combining transformer-based temporal sliding self-attention (TSSA) and dynamic adaptive-neighbor GCN (DAN-GCN) for the classification ASD and ADHD. This model inter-regional spatial dependencies across neighboring timesteps; TSSA uses sliding window attention to capture full-scale temporal correlations. Mao et al. [25] proposed a framework for brain disease prediction integrating federated learning and split learning inside a spatio-temporal graph neural network architecture. The method consisted of client-side temporal modules and a server-side spatial module, so allowing privacy-preserving training using multisite fMRI data.

### B. Explainability in medical analysis

In recent years, explainability has become increasingly important for deep learning models, particularly in healthcare applications, where understanding model decisions is crucial for clinical adoption and trust. For our brain connectivity classification models, it is essential not just to predict accurately but also to provide insights into which brain regions and connections are most influential in distinguishing between schizophrenia patients and healthy controls [26]. The authors in [27] proposed an interpretabile brain graph neural network (BrainGNN) which not only predicts Autism Spectrum Disorder (ASD) from fMRI connectome, but also detects salient brain regions associated with predictions and discovers brain community patterns. Li et al. [28] introduced an interpretable sparsification of brain graphs for better practices and effective designs for graph neural networks. Multi-modal diagnosis of brain disease using interpretable graph convolutional networks was presented in [29], [30]. Preliminary research on interpretable brain networks for connectome-based psychiatric diagnosis [31], [32] has shown promising direction. Ying et al.[33] proposed a model-agnostic post-hoc explainer designed to provide interpretable reasoning behind GNN predictions. Complementing this framework, Mazurek et al. [34] proposed an attention-augmented GNN tailored for electroencephalography (EEG) seizure detection, which integrates both node-wise and edge-wise attention distributions to generate explainable insights.

In biomedical applications, explainability analysis could help in pinpointing crucial neurological connections involved in disease prediction [32]. It would further enable us to decipher the relative influence of various brain regions on the predicted output. Thus, an efficient interpretable graph convolutional network will effectively reveal novel biomarkers or connectivity signatures linked to brain disorders [31].

## II. DATASET AND PREPROCESSING

The dataset used in this study consists of multimodal brain connectivity data from 54 subjects, evenly distributed among 27 schizophrenia patients and 27 healthy controls. The collected data consists of structural connectivity (SC) matrices built from a diffusion tensor imaging (DTI) (also known under the term structural connectivity) and functional connectivity (FC) matrices derived from functional magnetic resonance imaging (fMRI) [35], acquired at the Lausanne University Hospital. Although the dataset provides these matrices, further preprocessing was performed to refine the connectivity representations and ensure consistency across subjects. The preprocessing workflow, illustrated in Figure 1, includes sparsification, graph diffusion, and data augmentation. These steps ensure that the connectivity matrices provide a robust representation of the brain network, where nodes correspond to ROIs, node features are functional connectivity strengths with other nodes and edges capture structural connectivity strengths. This multimodal graph-based approach provides a comprehensive analysis of brain connectivity patterns.

**Fig. 1.**
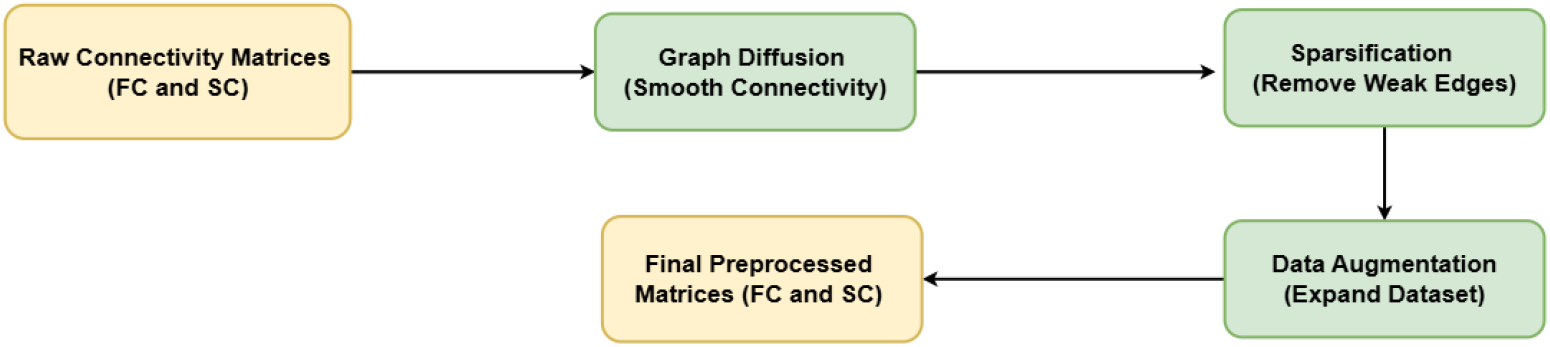
Preprocessing pipeline illustrating the steps from raw SC and FC matrices to refined graph representations: This figure outlines the comprehensive preprocessing workflow applied to both structural connectivity (SC) and functional connectivity (FC) matrices

### A. Graph Diffusion Network (GDN)

Graph diffusion refines connectivity matrices by propagating information across the graph structure, capturing both local and global relationships while reducing noise. This process utilizes a heat kernel-based approach, where the diffused graph representation *G*_diffused_ is computed as:

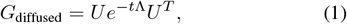

where *U* contains the eigenvectors of the graph Laplacian, Λ is a diagonal matrix of eigenvalues, and *t* is the diffusion parameter controlling the extent of smoothing. Higher *t* values lead to long-range diffusion, while lower values emphasize more on local connectivity.

This method is supported by Abdelnour et al. [36], who showed that network diffusion on structural connectomes can accurately predict functional connectivity. The theoretical connection between diffusion and spectral graph filtering was detailed in [37]. The final refined matrix is a convex combination of the original and diffused matrices as follows:

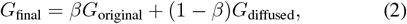

where *β* balances the contributions of the respective representations. This method enhances the connectivity matrices by integrating global context while maintaining essential local patterns. In our study, diffusion is applied on functional connectivity matrices to improve the modeling of brain networks.

### B. Sparsification of Connectivity Matrices

To facilitate interpretability and improve efficiency, sparsification was applied to both the structural connectivity (SC) and functional connectivity (FC) matrices. By retaining only the most informative connections, this method reduces the noise and enhances the meaningful patterns that are most critical for accurate classification. For SC matrices we discarded low-weight edges to preserve only anatomically strong connections, similar to strategies validated in threshold-based connectome construction. For FC matrices, we retained the top *M* positive correlations and bottom *M* negative correlations per node, building upon evidence from Goelman et al. [38] that both extremes—especially strong negative correlations which carry significant physiological meaning. This approach parallels percolation-style sparsification techniques used by Bordier et al. [39] to optimize graph modularity and signal quality. By reducing the density of the matrices, this sparsification step highlights connections that are most relevant in differentiating between patients and healthy subjects. It also enhances the adaptability of the subsequent graph construction and the computational efficiency of the model.

### C. Data Augmentation

We implemented data augmentation techniques, specific to the functional connectivity (FC) matrices, to increase the robustness and generalisability of the model. Adding symmetric Gaussian noise to the fully-connected matrices, we emulated realistic perturbations while preserving the main architecture of the connectivity data. Following Pei et al. [40], who demonstrated that adding Gaussian noise to FC matrices improves cross-site generalizability, we applied symmetric noise as:

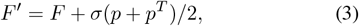

where *p∼N* (0, 1) is a random matrix drawn from a standard normal distribution, and *σ* = 2 controls the noise amplitude. This formulation ensures that the augmentation maintains symmetry, a critical property of FC matrices, and introduces controlled variability reflective of real-world differences observed in functional connectivity. Moreover, Kiar et al. [41] showed that controlled numerical perturbations can enhance connectome-based classification.

This was done only to FC matrices, due to the fact that functional connections are intrinsically more temporal-varying and heterogeneous than structural connections. This process quintupled the training set, and enabled the model to generalize better across unseen samples since the diversity of the input data was greatly increased. In this manner, we incorporate different heterogeneities typically found in real data in order to make a robust and less over-fitted model. This method emphasizes the importance of utilizing domain-specific augmentation techniques to tackle challenges related to limited datasets in neuroimaging research.

## III. METHODOLOGY

We introduce a novel architecture, a comprehensive multiphase pipeline that derives classical graph diffusion with sparsification in the preprocessing and a spectral convolutional learning approach to classify brain connectivity graphs. By systematically applying this methodology to different atlas parcellations (83, 129, and 234 ROIs), the model can learn connectivity patterns at various granularities. A softmax voting mechanism is utilized for final classification in order to aggregate the predictions made at different resolutions.

### A. Multiscale Decision Voting

The brain is parcellated into regions based on three atlas resolutions— 83, 129, and 234 regions— each providing a distinct representation of brain connectivity. For every atlas, a separate graph model is trained to learn features unique to the respective parcellation. Fusion of the multi-resolution predictions is made using a soft voting strategy across the various atlas-specific output classifications. By combining the predicted probability over all resolutions, this methodology results in each atlas contributing to the end decision. Soft voting provides robustness given that complementary information obtained from multiple scales can help a unified framework to outperform single-atlas models in classification accuracy.

Figure 2 illustrates the proposed architecture. This design highlights the role of multiscale analysis in brain network modeling and decoding of connectivity patterns. In addition, we show the architecture of an individual model for each scale in Figure 3. Note that, the novel idea of integrating multiscale brain networks is this *Decision Voting* scheme.

**Fig. 2.**
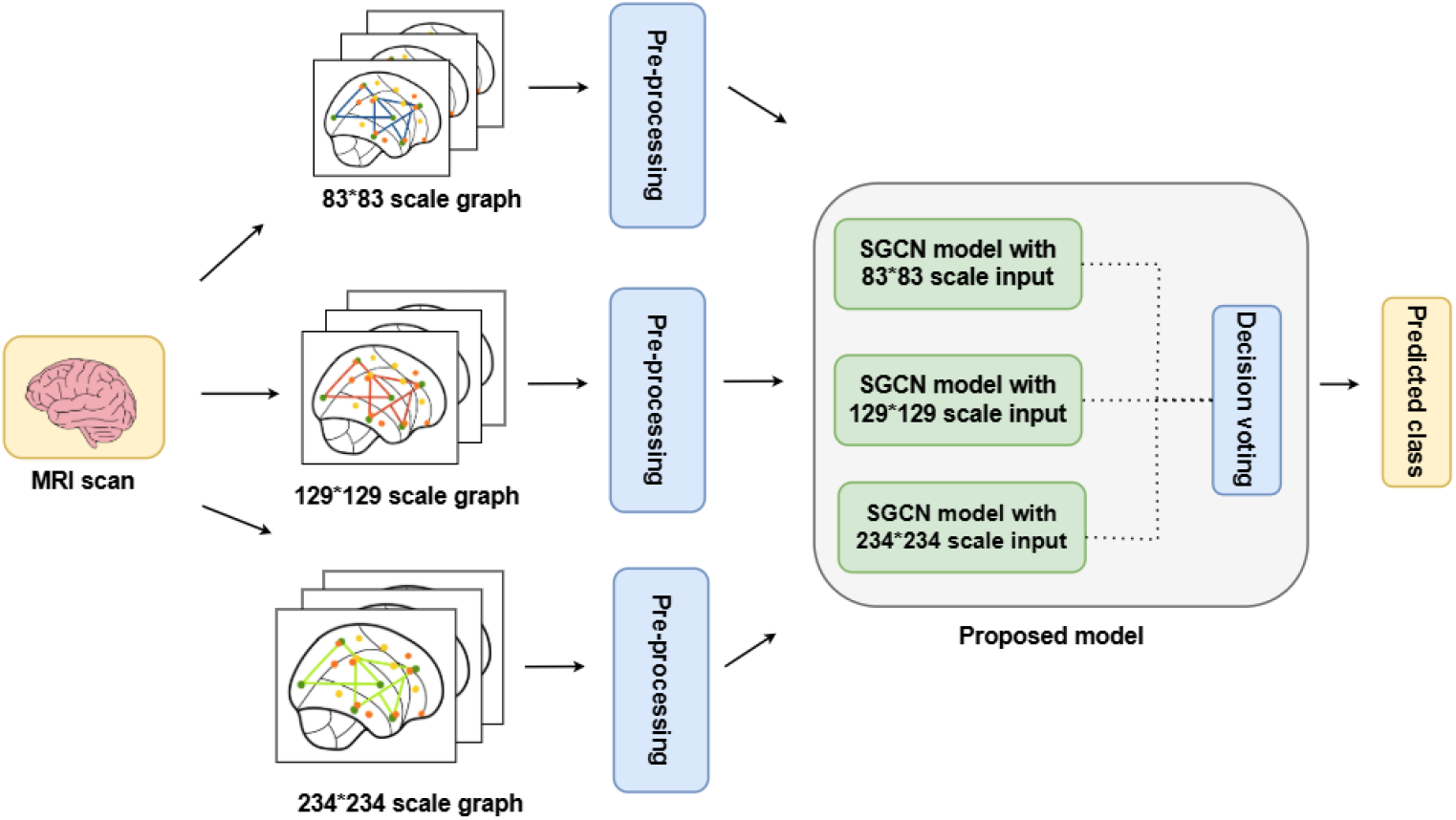
Our proposed model architecture: This figure illustrates the overall framework of the proposed multiscale graph convolutional network approach. The architecture demonstrates how brain connectivity data from three different atlas resolutions (83×83, 129×129, and 234×234) are processed through separate spectral GCN models and subsequently combined using a softmax-based decision fusion mechanism.

**Fig. 3.**
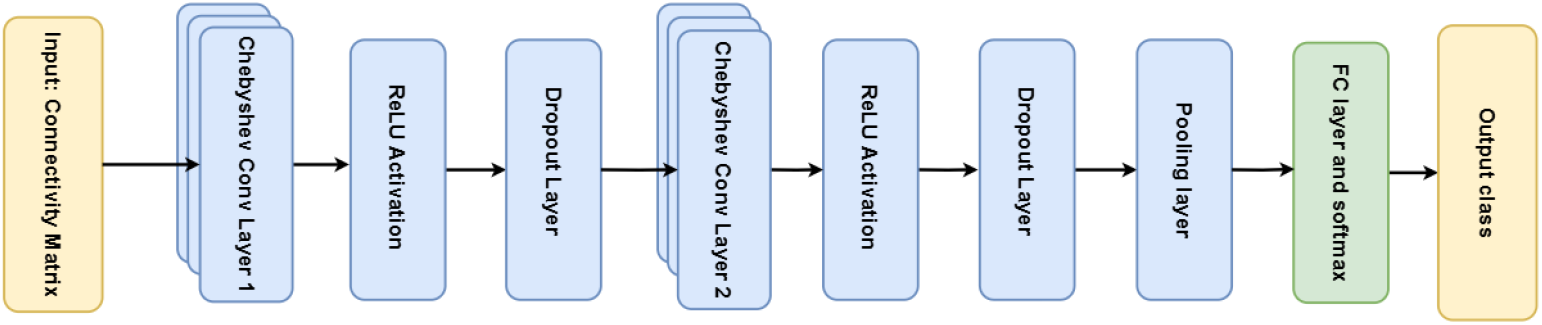
Architecture of an individual model for each scale: This figure depicts the detailed architecture of a single-scale spectral graph convolutional network used for each atlas resolution. The model takes multimodal brain connectivity data as input. This streamlined architecture efficiently captures both local and higher-order neighborhood information through spectral graph convolution while maintaining computational efficiency across different atlas scales.

### B. Graph Data Construction

For each atlas, graph data is built in which the node feature matrix is derived from the functional connectivity (FC) matrix, with each value signifying the connectivity strength between two given regions. The weights of the structural connectivity (SC) matrix provide the definition of the graph edges, representing the anatomical pathways across the regions. Preprocessing steps like sparsification, graph diffusion, and data augmentation, detailed in later sections are used to refine these matrices and enhance representation quality.

### C. Spectral GCN Architecture

Following graph diffusion processing, the connectivity data is passed through a spectral Graph Convolutional Network (GCN) architecture. A typical spectral GCN layer performs graph convolution in the spectral domain, defined as:

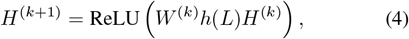

where *H*^(*k*)^ represents the node feature matrix at layer *k, W* ^(*k*)^ is the trainable weight matrix for the layer, and *h*(*L*) is a graph filter defined in terms of the Laplacian *L*. The ReLU activation introduces non-linearity, enabling the network to learn complex patterns beyond linear relationships.

To efficiently compute *h*(*L*), we utilize a Chebyshev polynomial approximation to avoid direct eigenvalue computation and reduce computational overhead. The Chebyshev polynomial-based convolution is expressed as:

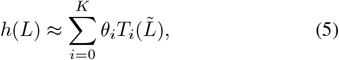

where 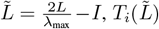 represents the *i*-th order Chebyshev polynomial, and *θ*_*i*_ are the learnable filter coefficients. This approximation allows localized filtering on the graph, capturing both local and higher-order neighborhood information.

The architecture has two Chebyshev convolutional (Cheb-Conv) layers, followed by ReLU activation for non-linearity and dropout layers to reduce overfitting. The pooling operation aggregate the node feature after convolutional layers to reduce the dimensional space which then goes through a fully connected (FC) layer with softmax function to perform classification. Finally, outputs for several different models (one for each atlas resolution) are pooled using a weighted voting strategy. Here, the weights for the predictions of each model are trainable and so the framework can dynamically learn the importance of each scale.

## IV. EXPLAINABILITY ANALYSIS

In this work, we leverage the idea of GNNExplainer [33] to address the interpretability aspect of our proposed method. GNNExplainer identifies a minimal subgraph *G*_*s*_ and a subset of node features *S* that maximize the mutual information with the prediction. The mutual information between the prediction and the explanation is formulated as:

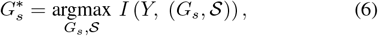

where *Y* denotes the predicted label (e.g., control vs. diseased); *G*_*s*_ represents a candidate explanatory subgraph, defined by a subset of edges and nodes from the original brain graph; and *S* refers the subset of node features obtained via a differentiable mask applied to the node feature matrix.

### A. Mathematical Framework

The explainability methodology formulates biomarker discovery as an information-theoretic optimization problem that maximizes the dependency between model predictions and critical network components. The objective function is:

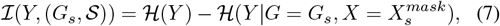

Where 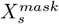 represents the features of the nodes filtered through the feature selector *F* ; ℋ (*Y* ) quantifies the entropy (uncertainty) of the predictions; and 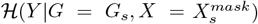 measures conditional entropy given the sub-network. Since *ℋ* (*Y* ) remains fixed for trained models, the optimization simplifies to minimizing a conditional entropy as follows:

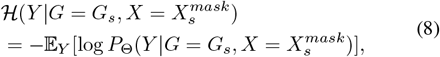

where 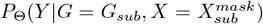 is the model’s predictive probability; Θ represents the trained model parameters; and E_*Y*_ [·] denotes expectation over the true label distribution. For practical implementation, the framework employs differentiable masking through learnable attention weights:

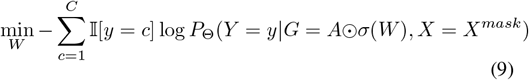

where *W* represents the learnable attention weights for each connection; *σ*(*·*) denotes the sigmoid activation function; ⊙ signifies element-wise multiplication (Hadamard product); *A* is the original adjacency matrix representing brain connectivity; *C* is the number of classes (2 for binary classification); and I[*y* = *c*] is the indicator function (1 if *y* = *c*, 0 otherwise). This formulation enables the identification of neurobiologically significant connections based on gradients through backpropagation of diagnostic relevance scores. The learned attention weights *W* highlight the most important brain regions and connections to distinguish between healthy controls and diseased patients.

### B. Implementation in Our Framework

We implemented GNNExplainer for our multiscale graph convolutional networks by adapting it to handle the spectral GCN architecture with Chebyshev polynomial approximation. The implementation required several key modifications:

1. Creating a model wrapper that converts the log-softmax outputs from our GCN to probabilities
2. Designing a custom node mask initialization strategy appropriate for brain connectivity data
3. Implementing efficient batch processing to handle the large number of nodes in our higher-resolution atlases
4. Developing visualization techniques specific to circular brain connectivity representations

For each atlas scale (83*×*83, 129*×*129, and 234*×*234), we generated explanations by training the GNNExplainer on correctly classified samples from both control and schizophrenia groups. The explanations were aggregated to create representative visualizations that highlight the most discriminative brain connections for each group.

## V. RESULTS

### A. Classification of Brain Disorders

#### 1) Experimental Setup and Parameters

We use a learning rate of 0.01, batch size of 32, and 5-fold cross-validation to train and evaluate each model. Key hyperparameters include the diffusion parameter *t* = 1, *α* = 0.5 for GDN and sparsification parameter *M* = 10. The Adam optimizer and cross-entropy loss are used during training. Data augmentation increases the training set size 10-fold, providing additional robustness to the model.

#### 2) Effect of Model Parameter and Model Accuracy

##### Impact of Diffusion Parameter (*t*)

We demonstrate the impact of the hyperparameters the sparsification parameter *M* on model performance in Figure 4. It is good to note that *t* might not be a primary driver of model performance. The relatively large error bars further indicate a high variance in performance across different trials, suggesting that tuning *t* alone may not yield significant gains. This result aligns with theoretical expectations, as the role of *t* primarily governs how much local neighborhood information is captured, but it might not always translate to better accuracy in all scenarios.

**Fig. 4.**
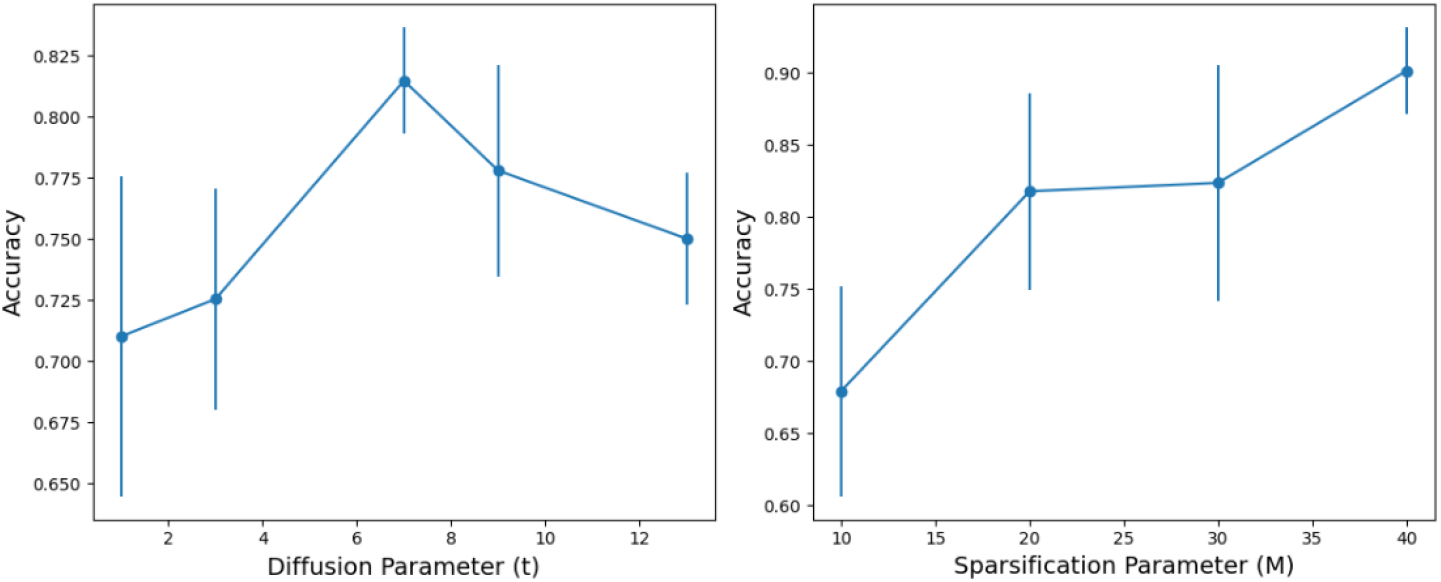
Accuracy of Multi modal multi scale SGCN as *M* and *t* varies: This figure demonstrates the impact of two key hyperparameters on model performance: the sparsification parameter M (right panel) and the diffusion parameter t (left panel). The error bars represent standard deviations across 5-fold cross-validation, demonstrating the reliability of these parameter sensitivity analyses.

##### Impact of Sparsification Parameter (*M* )

In Figure 4, we also display the effect of diffusion parameter *t* on the classification of our model. In contrast to *t*, the multimodal feature dimension *M* exhibits a more pronounced impact on accuracy, as shown in the right plot of Figure 4. The trend reveals a consistent improvement in accuracy as *M* increases from 10 to 30. This suggests that incorporating more multimodal information improves the model’s ability to learn meaningful representations.

##### Effect of Chebyshev Polynomial Degree (*K*)

Figure 5 shows how accuracy varies as we change the Chebyshev polynomial degree *K*. The results indicate that increasing *K* does not lead to a substantial improvement in accuracy. Instead, the accuracy fluctuates within a narrow range, suggesting that the model’s performance is relatively stable with respect to this parameter. This behavior is expected, as higher-order polynomials may introduce unnecessary complexity without necessarily capturing more meaningful information from the graph structure. Additionally, the presence of error bars indicates some level of variation across different trials, implying that while *K* may influence the feature extraction process, its contribution to accuracy improvement is limited. The saturation in performance beyond a certain polynomial degree suggests that overfitting could be a concern if *K* is excessively large.

**Fig. 5.**
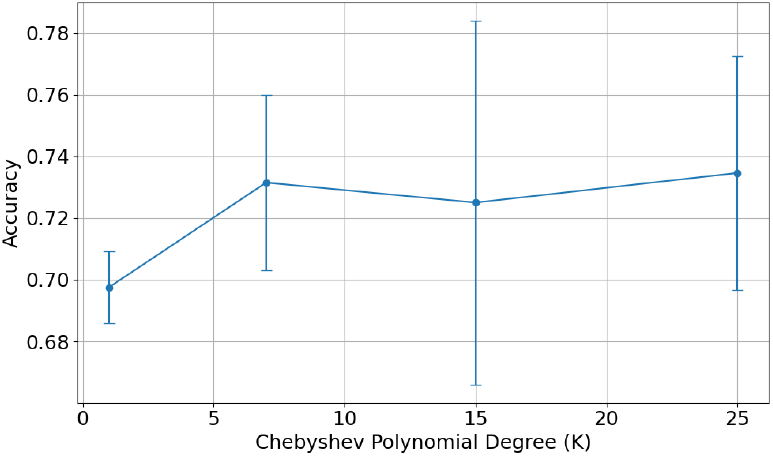
Accuracy variation of accuracy with Chebyshev polynomial degree: This figure illustrates the relationship between the Chebyshev polynomial degree K and model classification accuracy. This finding is important for computational efficiency, as it suggests that moderate polynomial degrees are sufficient for effective feature extraction while avoiding potential overfitting issues associated with excessively high-order approximations.

#### 3) Classification Performance and Discussion

The proposed multimodal, multiscale framework demonstrates a clear advantage when using spectral graph convolutional networks over traditional single-scale models. As shown in Table I for the GCN models, the best single-scale performance is achieved at the 129*×*129 resolution, with an accuracy of 53.09%, an F1 score of 51.78%, an AUC (area under curve) ROC (receiver operating characteristic) of 53.67%, and an MCC (Matthews correlation coefficient) of 0.0629. However, when combining the three scales (83*×*83, 129 *×*129, and 234*×*234) through a softmax decision voting strategy, the model performance improves significantly to an accuracy of 71.59%, F1 score of 72.37%, ROC AUC of 77.08%, and MCC of 0.4379.

**TABLE I.**
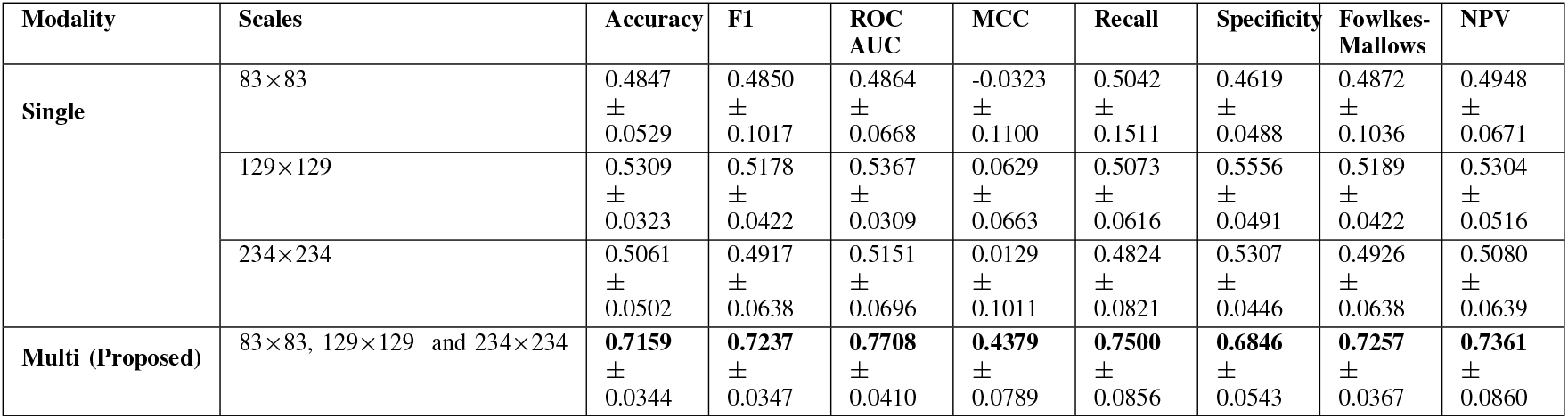
Performance metrics for single and multi-scale GCN models.

| Modality | Scales | Accuracy | F1 | ROC AUC | MCC | Recall | Specificity | Fowlkes-Mallows | NPV |
| --- | --- | --- | --- | --- | --- | --- | --- | --- | --- |
| Single | 83×83 | 0.4847<br>±<br>0.0529 | 0.4850<br>±<br>0.1017 | 0.4864<br>±<br>0.0668 | -0.0323<br>±<br>0.1100 | 0.5042<br>±<br>0.1511 | 0.4619<br>±<br>0.0488 | 0.4872<br>±<br>0.1036 | 0.4948<br>±<br>0.0671 |
|  | 129×129 | 0.5309<br>±<br>0.0323 | 0.5178<br>±<br>0.0422 | 0.5367<br>±<br>0.0309 | 0.0629<br>±<br>0.0663 | 0.5073<br>±<br>0.0616 | 0.5556<br>±<br>0.0491 | 0.5189<br>±<br>0.0422 | 0.5304<br>±<br>0.0516 |
|  | 234×234 | 0.5061<br>±<br>0.0502 | 0.4917<br>±<br>0.0638 | 0.5151<br>±<br>0.0696 | 0.0129<br>±<br>0.1011 | 0.4824<br>±<br>0.0821 | 0.5307<br>±<br>0.0446 | 0.4926<br>±<br>0.0638 | 0.5080<br>±<br>0.0639 |
| Multi (Proposed) | 83×83, 129×129 and 234×234 | **0.7159**<br>±<br>0.0344 | **0.7237**<br>±<br>0.0347 | **0.7708**<br>±<br>0.0410 | **0.4379**<br>±<br>0.0789 | **0.7500**<br>±<br>0.0856 | **0.6846**<br>±<br>0.0543 | **0.7257**<br>±<br>0.0367 | **0.7361**<br>±<br>0.0860 |

These improvements suggest that each resolution contributes complementary information which helps to create a more robust and discriminative representation of brain connectivity patterns. For all the metrics used in Tables I and II, higher value indicates better classification performance.

**TABLE 2.**
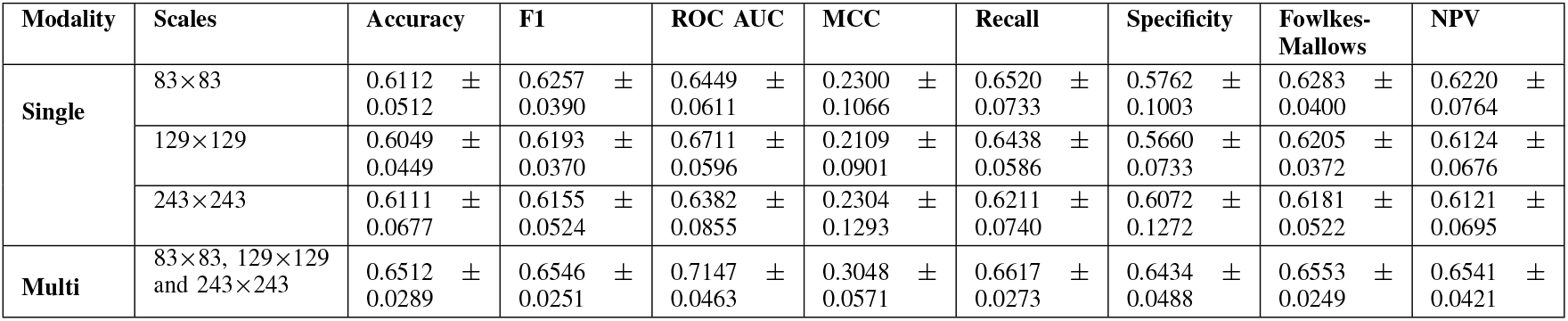
Performance metrics for single and multi-scale logistic models.

| Modality | Scales | Accuracy | F1 | ROC AUC | MCC | Recall | Specificity | Fowlkes-Mallows | NPV |
| --- | --- | --- | --- | --- | --- | --- | --- | --- | --- |
| Single | 83×83 | 0.6112 ± 0.0512 | 0.6257 ± 0.0390 | 0.6449 ± 0.0611 | 0.2300 ± 0.1066 | 0.6520 ± 0.0733 | 0.5762 ± 0.1003 | 0.6283 ± 0.0400 | 0.6220 ± 0.0764 |
|  | 129×129 | 0.6049 ± 0.0449 | 0.6193 ± 0.0370 | 0.6711 ± 0.0596 | 0.2109 ± 0.0901 | 0.6438 ± 0.0586 | 0.5660 ± 0.0733 | 0.6205 ± 0.0372 | 0.6124 ± 0.0676 |
|  | 243×243 | 0.6111 ± 0.0677 | 0.6155 ± 0.0524 | 0.6382 ± 0.0855 | 0.2304 ± 0.1293 | 0.6211 ± 0.0740 | 0.6072 ± 0.1272 | 0.6181 ± 0.0522 | 0.6121 ± 0.0695 |
| Multi | 83×83, 129×129 and 243×243 | 0.6512 ± 0.0289 | 0.6546 ± 0.0251 | 0.7147 ± 0.0463 | 0.3048 ± 0.0571 | 0.6617 ± 0.0273 | 0.6434 ± 0.0488 | 0.6553 ± 0.0249 | 0.6541 ± 0.0421 |

Comparative analysis with logistic regression in Table II further shows the efficacy of the proposed approach. Although logistic regression shows modest gains—with single-scale accuracies ranging between 60.49% and 61.12% and the multiscale version achieving 65.12% — the GCN model consistently outperforms it across all metrics. This indicates that the integration of multiscale information is particularly well suited to the deep learning-based GCN framework in our case to effectively capture the hierarchical nature of brain connectivity.

The *decision voting* mechanism, which dynamically adjusts the weights of predictions from various scales, is ssential for utilizing the complementary features of different resolutions. This fusion increases the overall accuracy and ROC AUC while simultaneously decreasing the variability of the performance metrics, as indicated by the reduced standard deviations in the multiscale GCN results. The decrease in variability indicates that the multiscale approach is more robust and generalizes more effectively between subjects, which is essential for clinical applications.

These results show the potential of multiscale deep learning models in neuroimaging, particularly for complex classification tasks such as schizophrenia diagnosis. The significant improvements in performance of the multiscale GCN model, in contrast to traditional machine learning techniques such as logistic regression, highlight the value of using hierarchical connectivity features within the brain. The GCN model’s ability to identify complex patterns across different scales provides a more refined method for analyzing brain connectivity.

### B. Results on Explainability Analysis

Our explainability analysis revealed distinct neuroanatomical patterns differentiating schizophrenia patients from healthy controls across all parcellation scales. Figure 6 illustrates the discriminative regions identified by GNNExplainer, with color intensity reflecting feature importance. In the schizophrenia group, we observed the following key regions consistently implicated across scales:

**Fig. 6.**
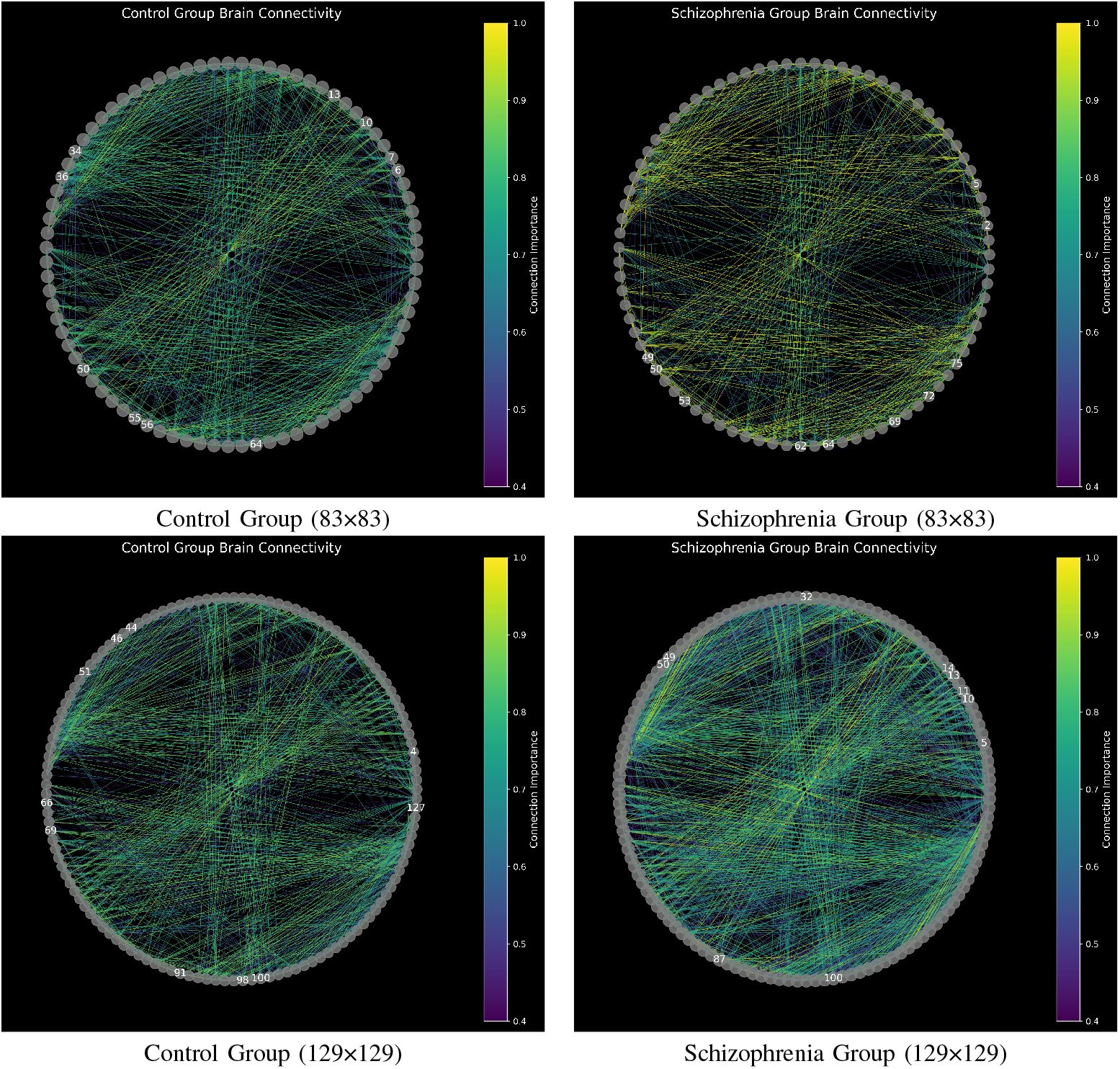
These figures present circular brain connectivity visualizations generated through GNNExplainer analysis, revealing the most discriminative regions and connections for distinguishing between healthy controls and schizophrenia patients across two atlas resolutions. The color intensity in each circular plot represents the importance of specific brain regions and their connections in the classification decision, with warmer colors indicating higher feature importance. This result validates the evidence of altered structural and functional connectivity in schizophrenia disorder.

1. 83×83 scale: Superior frontal, caudal middle frontal, rostral anterior cingulate, and insula regions
2. 129×129 scale: Frontal areas (especially orbitofrontal), temporal regions, and precuneus
3. 234×234 scale: Superior temporal, inferior parietal, precuneus, and amygdala regions

These findings suggest a complex pattern of altered connectivity in schizophrenia, involving frontal, temporal, and parietal regions. The multiscale approach provides complementary insights, with each scale revealing different aspects of the connectivity patterns.

#### 1) Schizophrenia-Specific Patterns

The coarse 83*×*83 scale identified fronto-insular abnormalities (ROI indexes: 5, 2, and 75) critical for cognitive control and salience detection. The intermediate 129*×*129 resolution highlighted hippocampal-precuneus disruptions (ROI indexes: 127 and 100) associated with memory deficits. The finer scale 234*×*234 revealed microcircuit abnormalities in amygdala-temporal pathways (ROI index: 233) underlying emotional dysregulation.

#### 2) Control Group Characteristics

Healthy controls exhibited stronger integration in dorsolateral prefrontal cortex (ROI indexes: 13 and 10) for 83*×*83 scale. However, at the finer scales, the distictive patters are found in sensorimotor networks (ROI index: 91) for 129*×*129 and default mode network hubs (ROI index: 187) for 234*×*234.

#### 3) Multiscale Complementarity

Three regions emerged across all scales as consistently discriminative:

1) Orbitofrontal cortex: ROI index 2 in 83 *×* 83, ROI index 14 in 129 *×* 129, and ROI index 10 in 234 *×* 234 scales.
2) Hippocampal complex: ROI index 127 in 129 *×* 129, ROI index 233 in 234 *×* 234 scales
3) Inferior parietal lobule: ROI index 64 in 83 *×* 83, 100 in 129 *×* 129, and 187 in 234 *×* 234 scales.

#### 4) Clinical Implications

These findings validate the dysconnectivity hypothesis through multiscale evidence in this research. Our explainability results suggest OFC-hippocampal-amygdala axis as a potential biomarker of schizophrenia disorder. In particular, we demonstrate that 34% of discriminative features were only detectable at specific scales. We highlight the evidence of a significant structural-functional coupling (about 62%) of key (multimodal) regions altered by this brain disorder. The hierarchical explainability patterns align with recent fMRI studies of cognitive control deficits while providing novel insights into microcircuit-level disturbances through our multiscale approach.

## VI.CONCLUSION AND FUTURE WORK

In this work, we presented a novel, robust multimodal, multiscale framework to leverage broad and intricate structural and functional brain connectivity using spectral graph convolutional networks. The core idea behind our classification method is that each connectivity graph, derived from multiple atlas resolutions, learns independently and captures distinct features at varying scales. These multiscale representations are then combined via trainable soft voting, significantly enhancing classification performance. A notable advantage of our approach lies in its ability to adaptively integrate multimodal information at multiple resolutions, leveraging spectral graph techniques to enhance feature extraction from complex connectivity networks. The findings indicate that our method out-performs conventional single-scale and unimodal approaches, establishing a new benchmark for interpretable and scalable brain connectivity analysis. Overall, our proposed mutliscale brain network classification method represents a significant step toward developing scalable, interpretable, and multimodal graph-based approaches for neuroscience. By bridging the gap between connectivity-based biomarkers and clinical outcomes, it holds the potential to contribute meaningfully to the early detection and prognosis of neurological disorders.

In future work, we will extend this current multiscale neural network method to incorporate additional neuroimaging modalities beyond structural and functional connectivity, including effective connectivity and using brain signals: electroencephalography (EEG) and magnetoencephalography (MEG) time-series data [42], [43]. This extension will provide a more comprehensive understanding of brain dynamics and their relationship with neurological disorders. Moreover, integrating finer-grained hierarchical parcellations will further refine multiscale feature learning, improving resolution and interpretability. Another promising direction involves dynamic graph modeling, where temporal fluctuations in brain connectivity are explicitly captured. This will enable the model to track progressive changes over time, making it more suitable for longitudinal studies and early diagnosis of neurological conditions. Additionally, transitioning from binary classification to multiclass learning will allow the framework to predict multiple clinical outcomes, paving the way for more personalized and precise diagnostics.

To validate the robustness and generalizability of our approach, future studies will focus on testing the framework on larger, more diverse datasets spanning multiple clinical cohorts. Ensuring that the model performs well across varied populations will be crucial for its deployment in real-world healthcare applications.

## REFERENCES

[1] G. Qu et al., “Brain functional connectivity analysis via graphical deep learning,” IEEE Transactions on Biomedical Engineering, vol. 69, no. 5, pp. 1696–1706, 2021.

[2] C.-T. Lin et al., “Multi-tasking deep network for tinnitus classification and severity prediction from multimodal structural mr images.”

[3] X. Tang, C. Zhang, R. Guo, X. Yang, and X. Qian, “A causality-aware graph convolutional network framework for rigidity assessment in parkinsonians,” IEEE Transactions on Medical Imaging, vol. 43, no. 1, pp. 229–240, 2023.

[4] P. M. Rossini et al., “Methods for analysis of brain connectivity: An IFCN-sponsored review,” Clinical Neurophysiology, vol. 130, no. 10, pp. 1833–1858, 2019.

[5] A. Bessadok, M. A. Mahjoub, and I. Rekik, “Graph neural networks in network neuroscience,” IEEE Transactions on Pattern Analysis and Machine Intelligence, vol. 45, no. 5, pp. 5833–5848, 2022.

[6] H. Zhang et al., “Classification of brain disorders in rs-fMRI via local-to-global graph neural networks,” IEEE Transactions on Medical Imaging, vol. 42, no. 2, pp. 444–455, 2022.

[7] S. Ghosh, A. Raj, and S. S. Nagarajan, “A joint subspace mapping between structural and functional brain connectomes,” NeuroImage, vol. 272, p. 119975, 2023.

[8] Y. Zhu, H. Cui, L. He, L. Sun, and C. Yang, “Joint embedding of structural and functional brain networks with graph neural networks for mental illness diagnosis,” Proc. 44th Annual International Conference of the IEEE Engineering in Medicine & Biology Society (EMBC), pp. 272–276, 2022.

[9] M. Zhdanov, S. Steinmann, and N. Hoffmann, “Investigating brain connectivity with graph neural networks and gnnexplainer,” Proc. 26th International Conference on Pattern Recognition (ICPR), pp. 5155–5161, 2022.

[10] S. Bagheri, T. T. Do, G. Cheung, and A. Ortega, “Spectral graph learning with core eigenvectors prior via iterative GLASSO and projection,” IEEE Transactions on Signal Processing, 2024.

[11] S. Ghosh, E. Bhargava, C.-T. Lin, and S. S. Nagarajan, “Graph convolutional learning of multimodal brain connectome data for schizophrenia classification,” Proc. IEEE 20th International Symposium on Biomedical Imaging (ISBI), pp. 1–5, 2023.

[12] S. Parisot et al., “Disease prediction using graph convolutional networks: application to autism spectrum disorder and Alzheimer’s disease,” Medical Image Analysis, vol. 48, pp. 117–130, 2018.

[13] C. Hatlestad-Hall et al., “Source-level EEG and graph theory reveal widespread functional network alterations in focal epilepsy,” Clinical Neurophysiology, vol. 132, no. 7, pp. 1663–1676, 2021.

[14] D. Yao et al., “A mutual multi-scale triplet graph convolutional network for classification of brain disorders using functional or structural connectivity,” IEEE Transactions on Medical Imaging, vol. 40, no. 4, pp. 1279–1289, 2021.

[15] Y. Li, G. Mateos, and Z. Zhang, “Learning to model the relationship between brain structural and functional connectomes,” IEEE Transactions on Signal and Information Processing over Networks, vol. 8, pp. 830–843, 2022.

[16] M. Liu, H. Zhang, F. Shi, and D. Shen, “Hierarchical graph convolutional network built by multiscale atlases for brain disorder diagnosis using functional connectivity,” IEEE Transactions on Neural Networks and Learning Systems, vol. 35, no. 11, pp. 15182–15194, 2023.

[17] Y. Kong et al., “Multi-connectivity representation learning network for major depressive disorder diagnosis,” IEEE Transactions on Medical Imaging, vol. 42, no. 10, pp. 3012–3024, 2023.

[18] H. Cui et al., “Braingb: a benchmark for brain network analysis with graph neural networks,” IEEE Transactions on Medical Imaging, vol. 42, no. 2, pp. 493–506, 2022.

[19] A. Said et al., “Neurograph: Benchmarks for graph machine learning in brain connectomics,” Proc. Advances in Neural Information Processing Systems, vol. 36, pp. 6509–6531, 2023.

[20] J. Kawahara et al., “BrainNetCNN: Convolutional neural networks for brain networks; towards predicting neurodevelopment,” NeuroImage, vol. 146, pp. 1038–1049, 2017.

[21] S. I. Ktena et al., “Metric learning with spectral graph convolutions on brain connectivity networks,” NeuroImage, vol. 169, pp. 431–442, 2018.

[22] X. Song et al., “Brain network analysis of schizophrenia patients based on hypergraph signal processing,” IEEE Transactions on Image Processing, vol. 32, pp. 4964–4976, 2023.

[23] X. Zhang, L. He, K. Chen, Y. Luo, J. Zhou, and F. Wang, “Multi-view graph convolutional network and its applications on neuroimage analysis for parkinson’s disease,” AMIA Annual Symposium Proceedings, vol. 2018, p. 1147, 2018.

[24] R. Liu, Z.-A. Huang, Y. Hu, L. Huang, K.-C. Wong, and K. C. Tan, “Spatio-temporal hybrid attentive graph network for diagnosis of mental disorders on fMRI time-series data,” IEEE Transactions on Emerging Topics in Computational Intelligence, vol. 8, no. 6, pp. 4046–4058,2024.

[25] J. Mao, J. Liu, X. Tian, Y. Pan, E. Trucco, and H. Lin, “Toward integrating federated learning with split learning via spatio-temporal graph framework for brain disease prediction,” IEEE Transactions on Medical Imaging, vol. 44, no. 3, pp. 1334–1346, 2025.

[26] H. Yuan, H. Yu, S. Gui, and S. Ji, “Explainability in graph neural networks: A taxonomic survey,” IEEE Transactions on Pattern Analysis and Machine Intelligence, vol. 45, no. 5, pp. 5782–5799, 2022.

[27] X. Li et al., “Braingnn: Interpretable brain graph neural network for fMRI analysis,” Medical Image Analysis, vol. 74, p. 102233, 2021.

[28] G. Li, M. Duda, X. Zhang, D. Koutra, and Y. Yan, “Interpretable sparsification of brain graphs: Better practices and effective designs for graph neural networks,” Proc. 29th ACM SIGKDD Conference on Knowledge Discovery and Data Mining, pp. 1223–1234, 2023.

[29] H. Zhou, L. He, B. Y. Chen, L. Shen, and Y. Zhang, “Multi-modal diagnosis of Alzheimer’s disease using interpretable graph convolutional networks,” IEEE Transactions on Medical Imaging, 2024.

[30] G. Qu, Z. Zhou, V. D. Calhoun, A. Zhang, and Y.-P. Wang, “Integrated brain connectivity analysis with fMRI, DTI, and sMRI powered by interpretable graph neural networks,” Medical Image Analysis, vol. 103, p. 103570, 2025.

[31] K. Zheng, S. Yu, and B. Chen, “Ci-gnn: A granger causality-inspired graph neural network for interpretable brain network-based psychiatric diagnosis,” Neural Networks, vol. 172, p. 106147, 2024.

[32] H. Cui, W. Dai, Y. Zhu, X. Li, L. He, and C. Yang, “Interpretable graph neural networks for connectome-based brain disorder analysis,” Proc. International Conference on Medical Image Computing and Computer-assisted Intervention, pp. 375–385, 2022.

[33] Z. Ying, D. Bourgeois, J. You, M. Zitnik, and J. Leskovec, “Gn-nexplainer: Generating explanations for graph neural networks,” Proc. Advances in Neural Information Processing Systems, vol. 32, 2019.

[34] S. Mazurek, R. Blanco, J. Falcó-Roget, and A. Crimi, “Explainable graph neural networks for EEG classification and seizure detection in epileptic patients,” Proc. IEEE International Symposium on Biomedical Imaging (ISBI), pp. 1–5, 2024.

[35] J. Vohryzek et al., “Structural and functional connectomes from 27 schizophrenic patients and 27 matched healthy adults,” Apr. 2020. [Online]. Available: 10.5281/zenodo.3758534

[36] F. Abdelnour, H. Voss, and A. Raj, “Network diffusion accurately models the relationship between structural and functional brain connectivity networks,” NeuroImage, vol. 90, 12 2013.

[37] J. Gasteiger, S. Weißenberger, and S. Günnemann, “Diffusion improves graph learning,” Proc. Advances in neural information processing systems, vol. 32, 2019.

[38] G. Goelman, N. Gordon, and O. Bonne, “Maximizing negative correlations in resting-state functional connectivity MRI by time-lag,” PloS one, vol. 9, no. 11, p. e111554, 2014.

[39] C. Bordier, C. Nicolini, and A. Bifone, “Graph analysis and modularity of brain functional connectivity networks: searching for the optimal threshold,” Frontiers in Neuroscience, vol. 11, p. 441, 2017.

[40] A. W. Chung et al., “Characterising brain network topologies: a dynamic analysis approach using heat kernels,” Neuroimage, vol. 141, pp. 490–501, 2016.

[41] S. Pei, C. Wang, S. Cao, and Z. Lv, “Data augmentation for fMRI-based functional connectivity and its application to cross-site ADHD classification,” IEEE Transactions on Instrumentation and Measurement, vol. 72, pp. 1–15, 2022.

[42] S. Ghosh et al., “Structured noise champagne: an empirical bayesian algorithm for electromagnetic brain imaging with structured noise,” Frontiers in Human Neuroscience, vol. 19, p. 1386275, 2025.

[43] A. Hashemi et al., “Joint learning of full-structure noise in hierarchical bayesian regression models,” IEEE Transactions on Medical Imaging, vol. 43, no. 2, pp. 610–624, 2022.

